# Sex-specific effects of gut microbiome on shaping bile acid metabolism

**DOI:** 10.1101/2024.06.27.601003

**Authors:** Yifei Yang, Liang Chi, Yun-Chung Hsiao, Kun Lu

## Abstract

Gut microbiome is a group of microorganisms that plays important roles in contributing to health and diseases. These bacterial compositions have been demonstrated to impact bile acids (BAs) profiles, either by directly metabolizing primary BAs to secondary BAs or indirect ways through host metabolism by influencing BAs synthesis, transportation and conjugation in liver. It has been observed sexually dimorphic gut microbiome and bile acids composition, with variations in expression levels of bile acid metabolizing genes in the liver. However, associations between sex-specific differences in gut microbiome and BAs profiles are not well understood. This study aimed to investigate whether gut microbiome could influence BAs profiles in host in a sex-specific manner. We transplanted cecum feces of male and female C57BL/6 mice to male mice and measured BAs concentrations in feces, serum and liver samples 7 days after fecal transplantation. We found different BAs profiles between mice with male and female gut microbiome, including altering levels and proportions of secondary BAs. We also observed varied expression levels of genes related to bile acid metabolism in the liver and distal ileum.Our results highlight sex-specific effects of gut microbiome on shaping bile acid metabolism through gut bacteria and regulation of host genes.

## 1 Introduction

Gut microbiome is a group of microorganisms, including the bacteria, commensal, and pathogens, residing in the gastrointestinal tract and plays a crucial role in contributing to health and diseases (Barko, McMichael, Swanson, & Williams, 2018; Oliphant & Allen-Vercoe, 2019; Shreiner, Kao, & Young, 2015). Imbalance of the normal gut microbiota is associated with inflammatory bowel disease (IBD), irritable bowel syndrome, obesity, and type 2 diabetes (Bull & Plummer, 2014). Gut microbiome has the ability to produce bioactive compounds that can mediate the activation of receptors in various cells, with its metabolic activity influencing overall host health (Holmes, Li, Marchesi, & Nicholson, 2012; Oliphant & Allen-Vercoe, 2019). It has been increasingly reported that gut microbiome-derived metabolites are key factors in the development of IBD, nonalcoholic fatty liver disease, and diabetes (Lavelle & Sokol, 2020; Puddu, Sanguineti, Montecucco, & Viviani, 2014; Wahlström, Sayin, Marschall, & Bäckhed,2016). One of the most important metabolites is BAs, which is produced by the host and further metabolized by gut microbiome (de Aguiar Vallim, Tarling, & Edwards, 2013). Recently, BAs have been regarded as important signaling molecules regulating glucose, lipid, and energy metabolism (Shapiro, Kolodziejczyk, Halstuch, & Elinav, 2018). It has been suggested that the potential involvement of sex□associated differences in gut microbiota and BAs in the unequal susceptibility and incidence of diseases (Razavi, Potts, Kelly, & Bazzano, 2019; L. Sheng et al., 2017). Previous studies have demonstrated gender-divergent of the gut microbiota composition and diversity and BAs concentrations (Fu, Csanaky, & Klaassen, 2012; Org et al., 2016). However, associations between sex-specific differences in gut microbiome and BAs profiles are not well understood.

BAs pools and compositions are regulated through bile acid synthesis, transportation and biotransformation in the liver and gut. Primary BAs are synthesized from cholesterol in liver and conjugated with taurine (predominant in mice) and glycine (predominant in humans) before secreting into bile. Then conjugated form of primary BAs is released into intestine, where BAs undergo deconjugation processes and biotransformation reactions to secondary BAs by gut microbiome. About 95% of BAs are reabsorbed in the ileum and returned to the liver through portal blood and the remaining 5% are excreted via feces. The process that BAs recycle between hepatocytes in the liver and enterocytes in the intestine, is called enterohepatic circulation.

Several factors can influence BAs profiles, including gut microbiome and host (Ridlon, Kang, Hylemon, & Bajaj, 2014). Gut microbiome appears to be a major regulator of the BAs in both direct and indirect ways. First, gut microbiome can directly regulate BAs composition by deconjugating conjugated primary BAs to unconjugated forms as well as metabolizing primary BAs to secondary BAs. Second, gut microbiome can also change BAs profiles in an indirect way through host metabolism. For example, after flowing back to liver through enterohepatic circulation, two major secondary BAs, including deoxycholic acid (DCA), lithocholic acid (LCA), serve as ligands of receptors, thus influencing BAs synthesis, transportation, and conjugation (Chiang, 2013; Dawson, Lan, & Rao, 2009; Li & Chiang, 2005). Different gut microbiome compositions may lead to different BAs profiles. It has been observed sexually dimorphic gut microbiome and BAs composition, with studies showing varying expression levels of BAs metabolizing genes in the liver (Durack & Lynch, 2019; Fransen et al., 2017; Fu et al., 2012; Org et al., 2016). However, specific ways of gut microbiome in different sexes influencing BAs remain unknown.

In this study, we will explore whether sex-specific gut microbiome could influence BAs profiles and the possible ways. Previous studies showed a dramatic difference in gut microbiome community structures between male and female mice (Chi et al., 2016; Council, 2000). To eliminate the influence of host sex, and genetics on bile acids profiles, male and female microbiome were used for transplantation.

## 2 Materials and Methods

### 2.1 Animal, antibiotics, fecal transplantation and sample collection

Female and male C57BL/6 conventional 7- to 8-week-old mice were purchased from Jackson Laboratories (Bar Harbor, ME) and housed for one week without any treatment to normalize gut microbiome. Mice were housed in sterilized microisolator cages and provided with ad libitum standard pelleted rodent diet and sterilized water. Workflow of this study is shown in Figure 1. Male C57BL/6 conventional mice were assigned to two groups: male mice transferred with male gut microbiome (M-M; n=9); and male mice transferred with female gut microbiome (F-M, n=9). Mice in both groups received a 3-day antibiotic treatment (cefoperazone, 0.5 mg/mL) through drinking water. Mice from M-M group were transplanted with mixed cecum feces from 3 C57BL/6 male donors; and mice from F-M group were transplanted with mixed cecum feces from 3 C57BL/6 female donors. All recipient mice had 7 days for bacteria colonization before conducting experiments. Body weight of recipients was measured at the start and endpoint (Supplementary Table 1), and no differences were observed between M-M and F-M groups (Supplementary Figure 2). Method of using CEF solution for ablation of bacteria has been confirmed by PCR as previously described (Chi et al., 2019). The concentration of the antibiotics was selected based on the previous literature (Sun et al., 2020). After a 3-day antibiotics treatment, mice feces were collected, and fecal DNA was extracted using a PowerSoil® DNA Isolation Kit (Qiagen) according to instructions of the manufacturer. DNA concentration and purity were measured using NanoDrop spectrophotometer (Thermo Fisher Scientific) to make sure the depletion of bacteria (Supplementary Table 2). The process of fecal transplantation experiments was based on method in previous paper (Suez et al., 2014). About 5 g of cecum feces were collected from donors and re-suspended in 5mL of PBS under anaerobic conditions, and centrifuged. 200 µl of the supernatant was used for gavage to receivers. To ensure a successful colonization of gut microbiome, fecal DNA was extracted, and DNA concentration was measured as above (Supplementary Table 3). Then, mice were euthanatized; heart blood, liver, and feces were collected and stored at -80□. All experiments were approved by the North Carolina Animal Care and Use Committee. Fresh mice liver tissue was snap-frozen in liquid nitrogen and then kept at -80°C. Fresh feces were collected into sterile fecal collection boxes and transported on dry ice before storage at −80□°C prior to further analysis.

**Figure 1.**
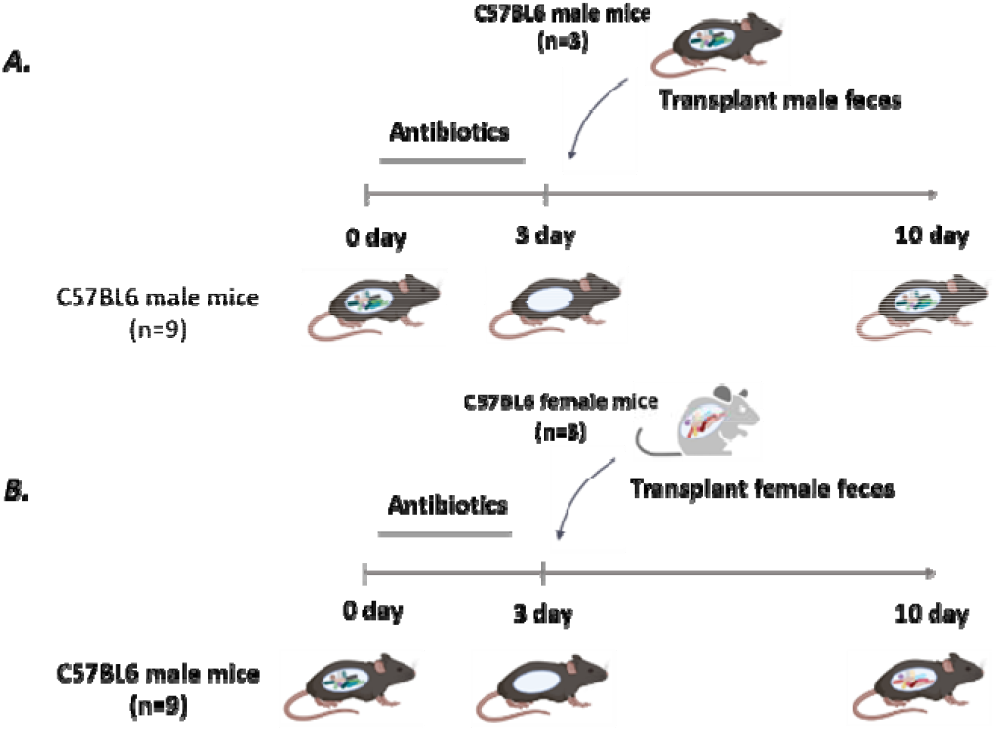
Experimental design. 18 C57BL6 male mice were divided randomly into two groups: (A). male mice transferred with male gut microbiome (M-M; n=9), (B). male mice transferred with female gut microbiome (F-M, n=9). After a 3-day cefoperazone (CEF) treatment, mice from M-M and F-M groups were transplanted with mixed cecum feces from 3 male and 3 female donors respectively, and had 7 days for bacteria colonization.

**Figure 2.**
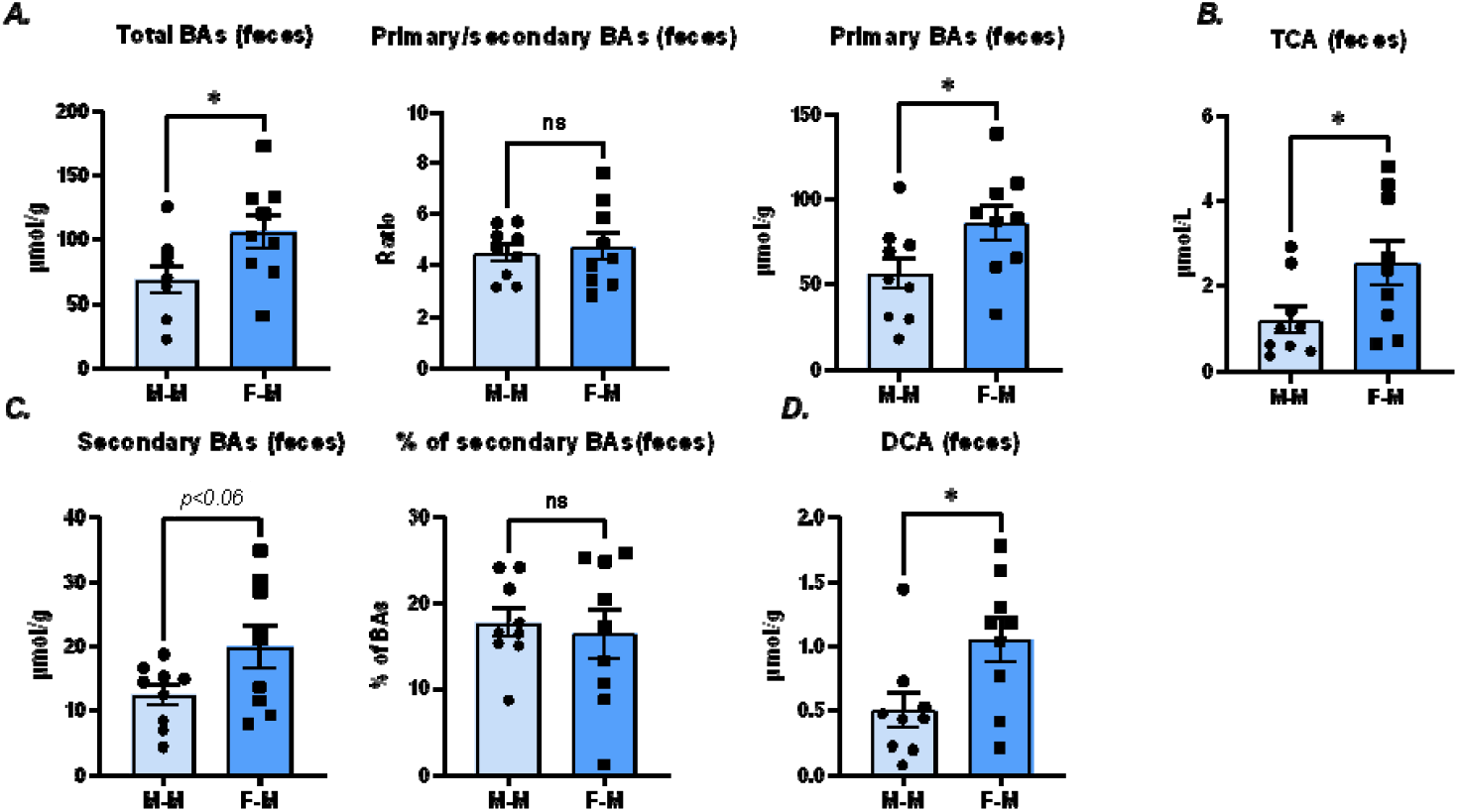
Female gut microbiome increases total BAs in feces. (A). Compared with male gut microbiome, mice with female gut microbiome were observed to significantly increase total and primary BAs levels without changing primary/ secondary BAs ratios. (B). Increasing primary BAs levels was mainly caused by increased levels of TCA. (C, D). Mice in F-M group had slightly increased secondary BAs levels, which was mainly caused by increased levels of DCA. (M-M: mice transplanted with male gut microbiome, F-M: mice transplanted with female gut microbiome; Data are expressed as mean ± SEM, unpaired t-test, ns: no significant difference;*p<0.05, **p<0.01, ***p<0.001; Error bar represents SEM; n = 9/group).

### 2.2 BAs measurement by mass spectrometry

The sample preparation was based on the method previously described with slight modifications (Alnouti, Csanaky, & Klaassen, 2008; Swann et al., 2011). For feces and liver samples, about 25 mg feces samples were weighed, added 20 µL of 1 µmol internal standards (IS), and extracted twice by 600 uL of ice-cold acetonitrile (ACN) and methanol (MeOH) in order. Samples were then homogenized using TissueLyzer at 50HZ for 15 min, centrifuged at 16,000 rpm for 10 min, and dried using SpeedVac (Thermo Fisher Scientific, Waltham, MA) for each extraction. Then the analytes were reconstituted with 200 uL MeOH and diluted to 1 mL water containing 0.2% formic acids (FA). The mixture was passed through an ISOLUTE C18 cartridge in the following order: 2 mL of MeOH, 2 mL of MeOH: water (0.1% FA) = (1:5), sample loading, 2 mL of MeOH: water (0.1% FA) = (1:5) and 1.2 mL MeOH. The products were dried using SpeedVac and reconstituted with 200 µL of MeOH: water (75: 25, v/v) for liquid chromatography-mass spectrometry analysis. For serum samples, 160 µL of ice-cold ACN was added to 20 µl serum-spiked with 20 µL of 1 µmol IS, vortexed, incubated at −20 °C for 30 min, and centrifuged at 15,000 rpm for 10 min. The supernatant was dried using SpeedVac and reconstituted in 100 µL of MeOH: H2O (75: 25, v/v) for liquid chromatography-mass spectrometry analysis. BAs were analyzed by liquid chromatography-electrospray ionization-triple quadrupole-mass spectrometry (LC-ESI-QQQ-MS) and operated in negative mode. The LC was carried out in an Agilent 1290 Infinity II UPLC system (Agilent Technologies, Santa Clara, CA) and MS was carried out by TSQ Quantis triple quadrupole mass spectrometry. Bile acid separation was achieved with an ACQUITY UPLC HSS T3 (2.1 mm × 100 mm, 1.8 µm) column (Waters Inc., Milford, MA, USA) with gradient elution at a flow rate of 0.4 ml/min and temperature maintained at 60 °C. Mobile phase A was composed of water containing 0.01% formic acid; mobile phase B was composed of acetonitrile containing 0.01% formic acid. The 26-min elution cycle started with 25% B, held for 2 min; then increased to 40% B in 13 min; then increased to 98% B at 20 min, held for 2 min; then decreased to 25% B at 22.5 min, and held for 3.5 min. The injection volume was 10 µL.

The following BAs were analyzed: tauro-α-muricholic acid (TαMCA), tauro-β-muricholic acid (TβMCA), tauro-ω-muricholic acid (TωMCA), tauroursodeoxycholic acid (TUDCA), taurochenodeoxycholic acid (TCDCA), taurodehydrocholic acid (TDHCA), taurodeoxycholic acid (TDCA), 3,7,12-trihydroxy-5-cholestanoic acid (THCA), taurocholic acid(TCA), cholic acid (CA), hyocholic acid (HCA), α-muricholic acid (αMCA), β-muricholic acid (βMCA), ω-muricholic acid (ωMCA), ursodeoxycholic acid (UDCA), murideoxycholic acid (MDCA), chenodeoxycholic acid (CDCA), dehydrocholic acid (DHCA), DCA, LCA, and glycocholic acid (GCA), glycohyocholic acid (GHCA), glycodeoxycholic acid (GDCA), glycodehydrocholic acid (GDHCA), glycolithocholic acid (GLCA), glycochenodeoxycholic acid (GCDCA), glycohyodeoxycholic acid (GHDCA), glycoursodeoxycholic acid (GUDCA). The quantification of BAs was based on the isotope dilution method with the responses of d4-TβMCA (for TαMCA, TβMCA, TωMCA), d4-TUDCA (for TUDCA), d9-TCDCA (for TCDCA), d6-TDCA (for TDHCA, TDCA), d4-CA (for THCA, TCA, CA, αMCA, βMCA, ωMCA), d4-UDCA (for UDCA, d9-CDCA (for MDCA, CDCA), d6-DCA (for DHCA, DCA), and d4-LCA (for LCA), d4-GCA (for GCA, GHCA), d6-GDCA(for GDCA,GDHCA), d4-GLCA(for GLCA), d4-GUDCA(for GCDCA, GHDCA,GUDCA). Multiple reaction monitoring transitions used for each bile acid were shown in Supplementary Table 4. At least eight-point calibration curves Analytical standards with concentrations ranging from 0.1-1000 nM were employed to generate least eight-point calibration curves with regression coefficients ≥ 0.99. Limits of detection for BAs were all below 0.5 nM.

### 2.3 Real time-quantitative PCR for gene expression measurement

The RNAs in liver and distal ileum tissues in RNA liter were isolated using RNeasy Mini Kit (Qiagen) according to the manufacturer ‘s instructions. The resultant RNA was estimated its concentration and purity by Nanodrop 2000 (Thermo Fisher Scientific) and digested genomic DNA contaminants using DNA-free™ DNA Removal Kit (Thermo Fisher Scientific). Then the DNA-digested RNA was used to synthesize cDNA using iScript™ Reverse Transcription Supermix (Bio-Rad Laboratories), and the products were diluted fivefold for quantitative real-time PCR (qPCR). qPCR experiments utilized Bio-Rad CFX96 Touch Real-Time PCR Detection System, and each reaction contained 10 μL SsoAdvancedTM Universal SYBR Green Supermix (Bio-Rad Laboratories), 1 μL 0.5 μM of each primer, and 5 μL template DNA. qPCR conditions started at a denaturation step at 95°C carried for 10 min followed by 35 PCR cycles that consisted of a denaturation (95°C for 10 s), annealing (54°C–55°C for 30 s), extension step (72°C for 30 s) and a final melting curve analysis of raising the temperature from 65–95°C in 0.5°C increments for 0.05 s each. All measurements were normalized by the expression of β-actin gene, which is considered as the housekeeping gene. Primer sequences and their annealing temperatures are listed in Supplementary Table 5. The relative expression of each gene was calculated using the ⍰ ⍰CT method with CFX (version 3.1) manager software (Bio-Rad).

### 2.4 Statistical analysis

Mice sample sizes were guided by previous animal studies in our laboratory. All data are presented as mean ± SEM. Statistical analysis was performed with Graph-Pad Prism 9 and the difference between groups was calculated by Student ‘s unpaired t-test. P < 0.05 was considered as a significant difference.

## Results

### 3.1 BAs profiles in feces

First, we investigated BAs profiles in feces, which are a useful proxy to study the gut microbiota. Globally, we observed significantly increasing levels of total and primary BAs in male mice transplanted with female gut microbiome group (F-M group), without changing ratios of primary to secondary BAs (Figure 2A). Higher levels of TCA, one of the most abundant primary BAs in mice, were observed in mice from F-M group (Figure 2B).

Compared with male mice transplanted with male gut microbiome group (M-M group), mice with female gut microbiome moderately increased secondary BAs levels, but this did not reach significance (Figure 2C). Levels DCA was observed significant increase. (Figure 2D). Gut microbiome in different sex didn’t affect unconjugated/conjugated BAs ratio (Supplementary Figure 1).

### 3.2 BAs profiles in serum

Next, we investigated serum BAs composition and found that total BAs levels as well as primary BAs levels were similar in two groups; ratios of primary to secondary BAs were lower in F-M group mice than in M-M group (Figure 3A). Mice in F-M group had a decreased primary BAs proportion without affecting the levels, and one of the major primary BAs in mice, CA, was observed a decreased proportion (Figure 3A and B).

**Figure 3.**
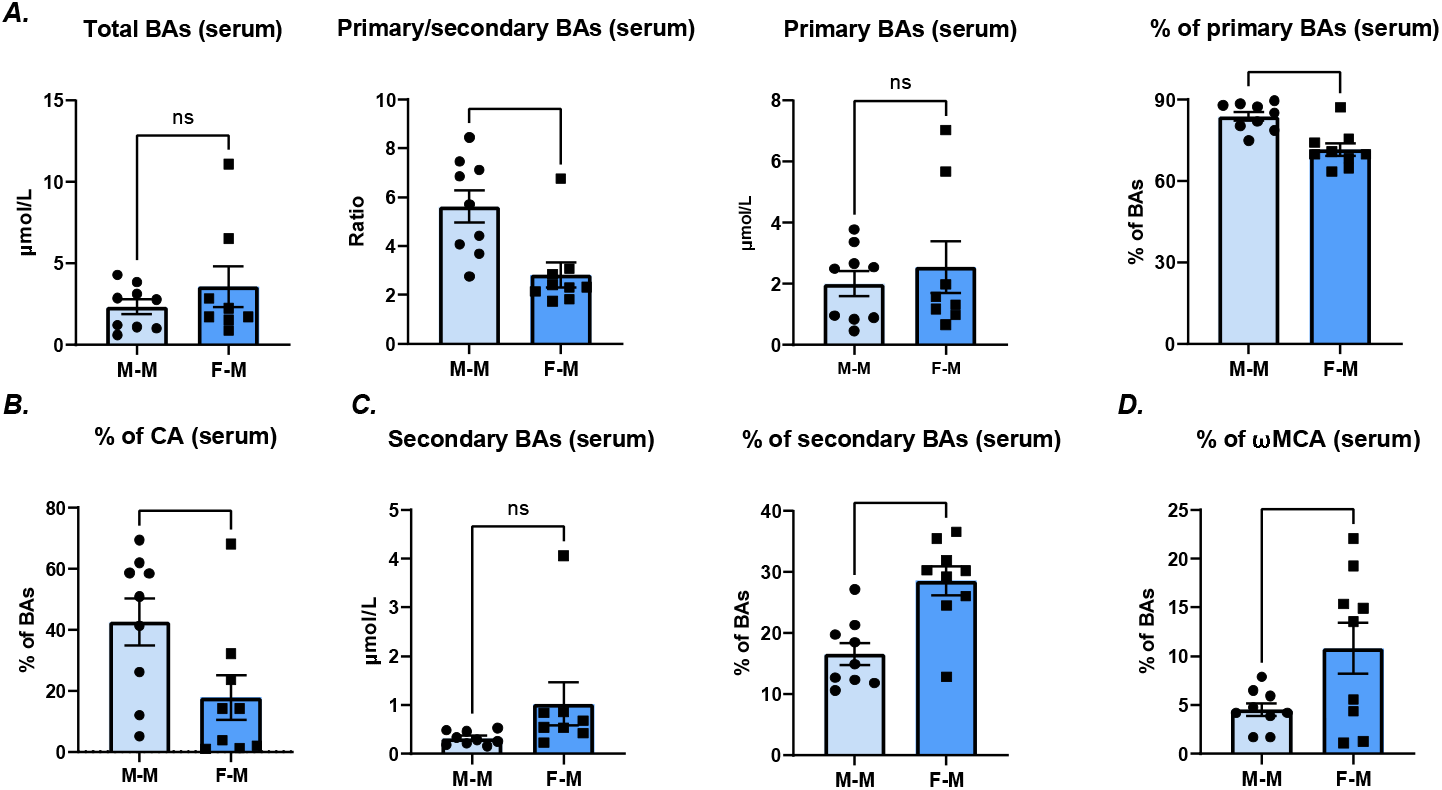
Female gut microbiome increases secondary BAs proportion in serum. (A, B). Female gut microbiome decreased primary BAs proportion without changing total and primary BAs levels, which was mainly caused by a decreased proportion of CA. (C, D). Female gut microbiome increased secondary BAs proportion rather than levels, which was mainly caused by an increased proportion of ωMCA. (M-M: mice transplanted with male gut microbiome, F-M: mice transplanted with female gut microbiome; Data are expressed as mean ± SEM, unpaired t-test, ns: no significant difference; *p<0.05, **p<0.01, ***p<0.001; Error bar represents SEM; n = 9/group).

Secondary BAs were found to have a higher percentage in mice of F-M group, but the levels of secondary BAs were constant in two groups (Figure 3C). One of the most abundant secondary BAs, ωMCA, was observed an increased proportion (Figure 3D). Like feces, gut microbiome of different sex didn’t affect unconjugated/conjugated BAs ratio (Supplementary Figure 1).

### 3.3 BAs profiles in liver

Finally, we investigated BAs composition in liver, where primary BAs are synthesized. Female gut microbiome led to unaffected total and primary BAs levels; however, it significantly decreased the ratio of primary to secondary BAs (Figure 4A). A lower proportion of TβMCA was shown in mice of F-M group (Figure 4B).

**Figure 4.**
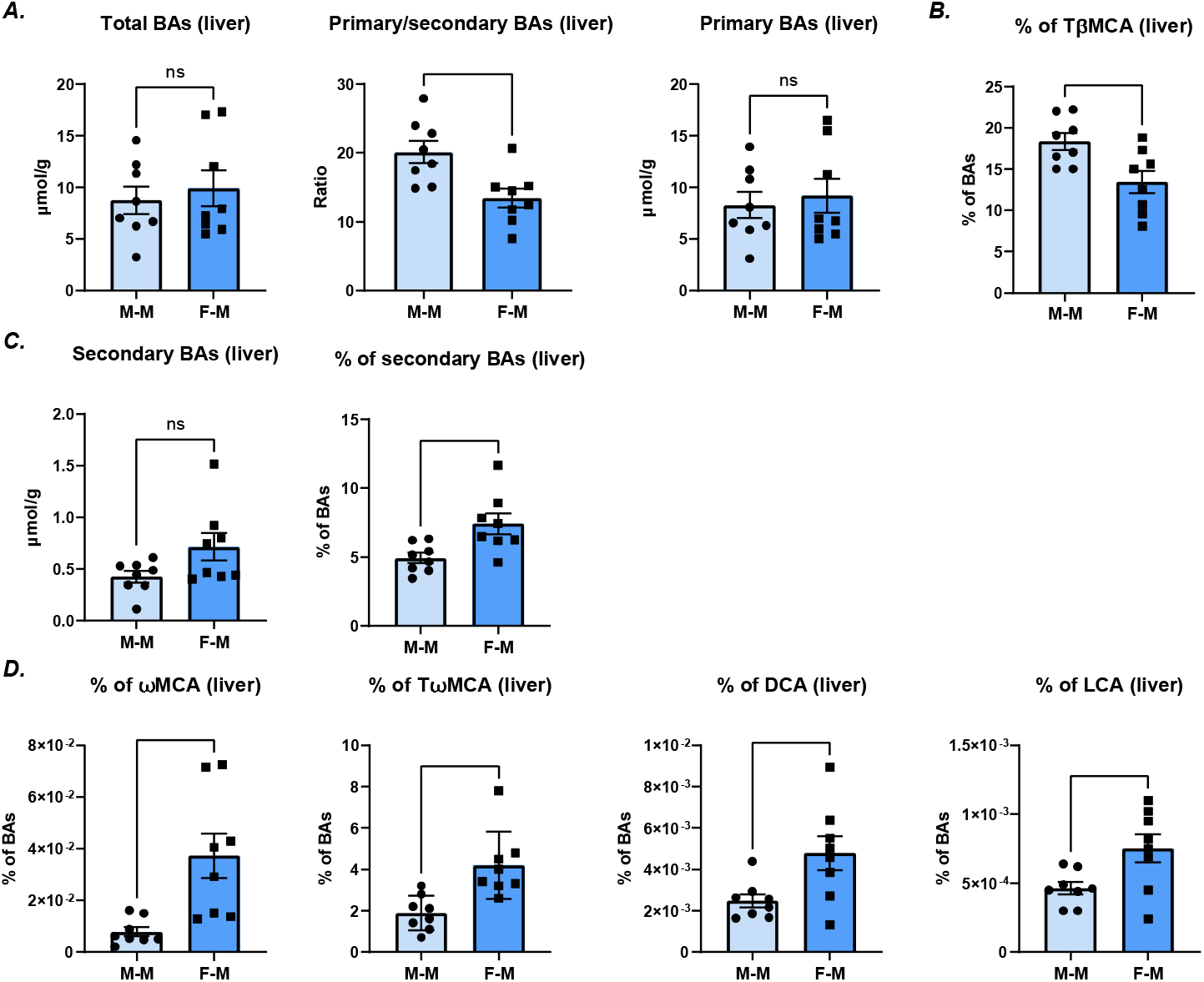
Female gut microbiome increases secondary BAs proportion in liver. (A, B). Female gut microbiome decreased primary BAs proportion without changing total and primary BAs levels, which mainly caused by a decreased proportion of TβMCA. (C, D). Female gut microbiome increased secondary BAs proportion rather than levels, which was caused by an increased proportion of major BAs, including ωMCA, TωMCA, DCA, and LCA. (M-M: mice transplanted with male gut microbiome, F-M: mice transplanted with female gut microbiome; Data are expressed as mean ± SEM, unpaired t-test, ns: no significant difference; *p<0.05, **p<0.01, ***p<0.001; Error bar represents SEM; n = 9/group).

A higher proportion of secondary BAs was observed in mice of F-M group without changing the levels (Figure 4C). Specific secondary bile acid species are shown in Figure 3D, where the relative abundance of ωMCA, TωMCA, DCA and LCA were significantly elevated in mice of F-M group. Gut microbiome of different sex didn’t affect unconjugated/conjugated BAs ratio (Supplementary Figure 1).

### 3.4 Transcript profiling in liver and distal ileum

To test whether gut microbiome can influence BAs through host metabolism, hepatic and distal ileum mRNA levels were quantified by qPCR. As shown in Figure 5A, the expression of rate-limiting enzyme CYP7A1 didn’t show differences between M-M and F-M groups.

**Figure 5.**
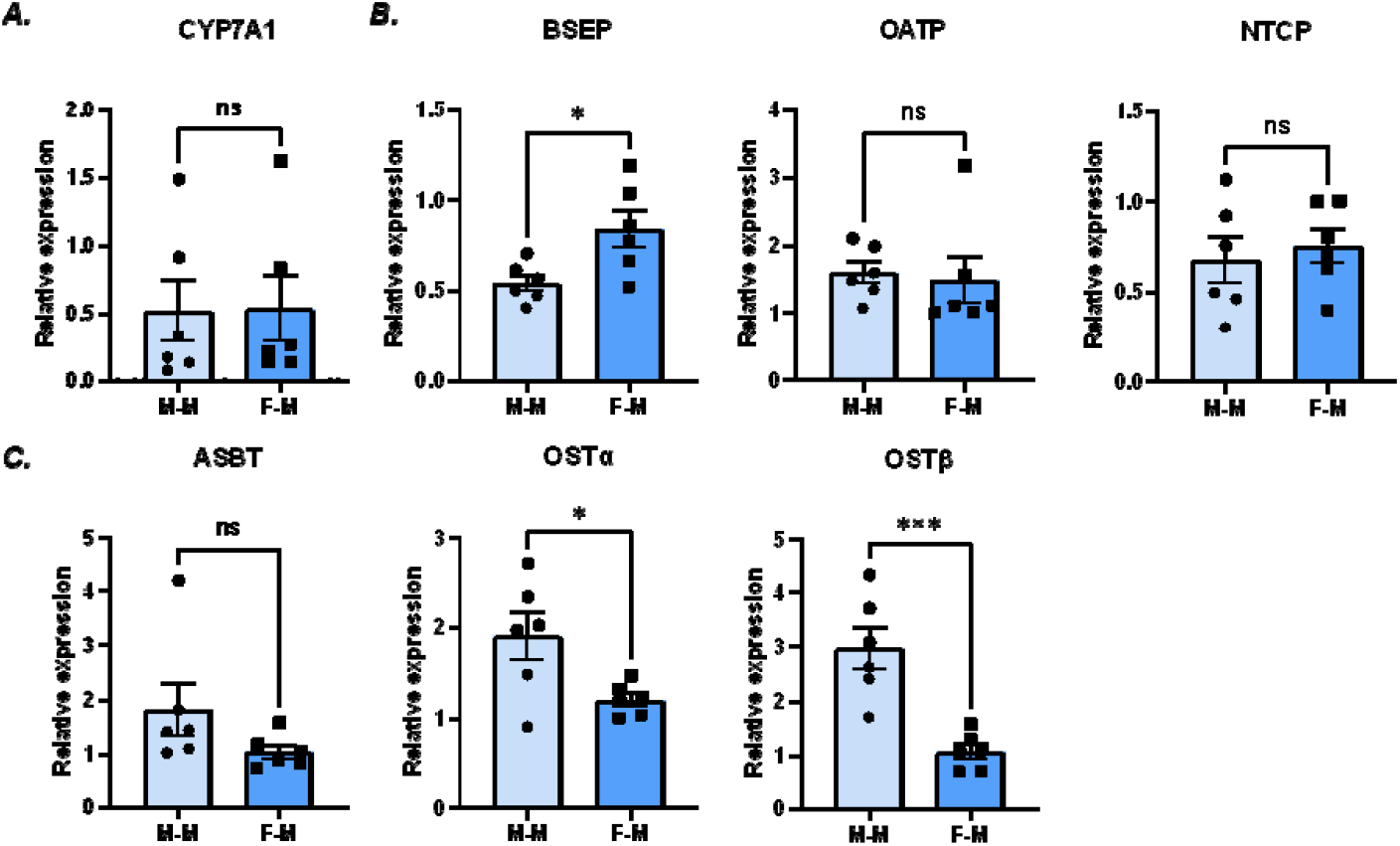
BAs transporters were regulated in liver and distal ileum. (A). BAs synthesis gene (CYP7A1) was not altered in two groups. (B). In liver, bile acid efflux transporter (BSEP) was significantly upregulated; uptake transporters (OATP and NTCP) were barely perturbed; (C). In distal ileum, uptake transporter (ASBT) showed no difference; efflux transporters (OSTα, OSTβ) significantly increased. (M-M: mice transplanted with male gut microbiome, F-M: mice transplanted with female gut microbiome; Data are expressed as meanL±LSEM, unpaired t-test, ns: no significant difference; *p<0.05, **p<0.01, ***p<0.001; n = 6 /group).

In liver, bile salt export pump (BSEP), was significantly highly expressed in mice with female gut microbiome than that of male gut microbiome; and two BAs uptake transporters, sodium taurocholate cotransporting polypeptide (NTCP) and organic anion transporting polypeptide (OATP), remained relatively constant (Figure 5B). BAs conjugation related gene, BAAT, had similar expression levels in two groups (Supplementary Figure 1). In distal ileum, bile acid uptake transporter, apical sodium dependent bile acid transporter (ASBT), showed no difference in two groups; two efflux transporters, organic solute transporter α and β (OSTα, OSTβ), significantly downregulated (Figure 5C).

## Discussion

The present study revealed that sex-specific gut microbiome influences BAs profiles in feces, serum, and liver (Figure 2-4). We observed a female-specific increase of secondary BAs levels in feces (Figure 2) along with significant elevations in secondary BAs proportions in serum and liver (Figure 3,4). These findings suggest the potential gut microbiome effects that sex-specific gut microbiome can directly influence BAs profiles, which female-predominant gut microbiome may have the ability to produce more secondary BAs. Two of the secondary BAs, ωMCA, DCA, exhibited predominate changes. ωMCA, a highly abundant secondary BA in mice, was observed to increase the percentage in serum and liver (Figure 3,4). DCA, one of the important secondary BAs, displayed increased levels in feces and elevated percentages in liver (Figure 2,4).

Gut microbiome is responsible for biotransformation reactions, metabolizing primary BAs (CA and CDCA) to secondary BAs (DCA and LCA, respectively) through BA 7α/β -dehydroxylation, and reactions of βMCA through BAs 6β-epimerization results in the formation of the secondary BAs, ωMCA (Eyssen, De Pauw, Stragier, & Verhulst, 1983; Wahlström et al., 2016). 7α/β - dehydroxylation bacteria have been characterized in diverse gut microbes. These are largely represented by species belonging to Clostridium spp., such as *C. scindens, C. hiranonis, C. hylemonae* (Clostridium cluster XIVa) and *C. sordelli* (Clostridium cluster XI) (Ridlon, Kang, & Hylemon, 2006). We assume that BAs biotransformation reactions, including 7α/β-dehydroxylation and 6β-epimerization activities, are different between male and female gut microbiome, thus leading to sex-specific changes in secondary BAs. Relative abundance of bacteria species that produce secondary BAs may have an expansion in female-transplanted mice. Enzymes encoded by bile acid inducible (bai) genes, which have been reported to be responsible for 7α/β-dehydroxylation reactions (Devlin & Fischbach, 2015), may have a higher amount in female gut microbiome than in male. Since CA can be converted to DCA, this might explain the decrease of the proportion of CA in serum and the increase of DCA in the liver and feces (Chiang & Ferrell, 2020). In the liver, we observed decreased proportions of TβMCA and increased proportions of ωMCA and TωMCA, this might be explained by the fact that gut bacteria convert β-MCA to ωMCA (Chiang & Ferrell, 2020).

Bile salt deconjugation is another important function carried out by gut microbiome enzyme bile salt hydrolase (BSH). In BAs metabolism, two processes are heavily studied about conjugation and deconjugation: hepatocyte conjugation reactions and gut bacteria deconjugation reactions. The expression levels of BAAT, the only BAs conjugation gene in liver (Pellicoro et al., 2007), were no different, which means that sex-specific gut microbiome may not alter BAs conjugation ability in liver. There were similar ratios of unconjugated and conjugated BAs in two groups (Supplementary Figure 1). Taken together, sex-specific effects of gut microbiome may have similar deconjugation activities between males and females.

BAs synthesis enzyme and transporters in the hepatocytes and enterocytes are listed in Figure 5. In the liver, primary BAs are produced from cholesterol and rate-limiting enzyme (CYP7A1) are responsible for the major pathway for bile acid synthesis. Then BAs are conjugated with taurine and glycine and secreted through BSEP in hepatocytes into bile. After BAs are released from the gallbladder to intestinal lumen, BAs are deconjugated (BSH) and metabolized (bai enzyme) by gut bacteria to form secondary BAs. Most BAs (about 95%) will be reabsorbed in intestine, with active transport by ASBT in the ileal enterocytes and passive transport in colon part (not shown in Figure 5) (Dawson & Karpen, 2015); the left 5% is excreted via feces. After uptake by ilea enterocytes, BAs are secreted into the portal circulation mediated by OSTα/β and then taken up by NTCP (predominate uptake of conjugated BAs) and OATP (predominate uptake of unconjugated BAs) in liver; only a small portion of BAs reaches systemic circulation. In Figure 5, several genes regulating bile acid metabolism in liver and distal ileum displayed more or less sex-specific expression patterns. Unchanged BAs synthesis gene (CYP7A1) expression revealed that sex-specific gut microbiome may not alter BAs synthesis (Figure 5A). Changed expression levels of transporter genes in liver and distal ileum suggested that sex-specific gut microbiome changes BAs profiles by influencing BAs transportation (Figure 5B and C). In F-M group, increased expression of efflux transporter (BSEP) in liver and unaltered expression levels of uptake transporter (ASBT) showed that more BAs excreted from hepatocytes, but unchanged levels of BAs were absorbed by enterocytes, leading to accumulation of total BAs excretion in feces. Significant lower expression levels of OSTα and OSTβ were observed in F-M group, indicating less BAs flowed into the portal circulation; expression levels of NTCP and OATP remained constant in two groups, which means similar levels of BAs were absorbed by liver. Therefore, it was speculated that more BAs reached to systemic circulation of mice in M-M group These results show that gut microbiome can indirectly change BAs profiles by changing its transportation rather than synthesis and conjugation.

BAs play an important role in host health. TCA has been reported to be associated with the promotion of germination of *Clostridioides difficile* and the progression of liver cirrhosis (Faintuch & Faintuch, 2019; Liu et al., 2018). A study demonstrated the protective effects of TCA in reducing hepatic lipid accumulation (Xu et al., 2022). It is reported that increased levels of DCA have been associated with obesity and colon cancer in mice (Bernstein, Bernstein, Payne, Dvorakova, & Garewal, 2005; Yoshimoto et al., 2013). LCA is found to be associated with a variety of hepatic and intestinal diseases (W. Sheng, Ji, & Zhang, 2022). Secondary BAs, including DCA and LCA, are shown to have detrimental effects of causing colon cancer by modulating M3R and Wnt/β-catenin signaling pathways (Farhana et al., 2016). Other researchers showed that secondary BAs can inhibit the growth of *Clostridioides difficile* (Thanissery, Zeng, Doyle, & Theriot, 2018). Recently, ωMCA has demonstrated to regulate liver cancer through decreasing expression of CXCL16 mRNA and natural killer T cell accumulation in liver (Ma et al., 2018).

BAs are ligands for many host cell receptors, including farnesoid X receptor (FXR), G protein-coupled receptor TGR5, pregnane-activated receptor (PXR), Vitamin D receptor (VDR), constitutive androstane receptor (CAR) (Ridlon et al., 2014; Wagner et al., 2005). Different BAs species can be served as agonists and antagonists to activate or inactivate the receptors, thus influencing the downstream signaling pathways including glucose, lipid, and energy homeostasis (Ding, Yang, Wang, & Huang, 2015; Thomas et al., 2009; Watanabe et al., 2006). Compared with primary BAs, secondary BAs seems to have greater potential to activate receptors; DCA and LCA are more efficacious ligands to activate TGR5, PXR, VDR compared with CDCA and CA (Hylemon et al., 2009; Kawamata et al., 2003; Ridlon et al., 2014; Studer et al., 2012). For FXR, CDCA is the most potent ligand, followed by LCA, DCA, and CA (Hylemon et al., 2009); while TβMCA has been identified as FXR antagonist (Sayin et al., 2013). FXR is an important nuclear receptor expressed in liver and intestine, which have a negative feedback control of BAs synthesis through regulating CYP7A1. We have observed changes of levels or proportions of CA, DCA, LCA, and TβMCA, which might activate or inactivate the receptors and their downstream pathways. FXR is also proven to regulate BAs transporters, including OSTα, OSTβ and BSEP (Ballatori et al., 2009; Claudel, Staels, & Kuipers, 2005). As shown in Figure 5, the three transporters we found to have significant changes between two groups. BAs transporters may be regulated by BAs through receptors (Ballatori et al., 2009; Claudel et al., 2005; Weerachayaphorn et al., 2012).

## Conclusion

In summary, our finding suggests that sex-specific gut microbiome may influence BAs metabolism by directly producing higher amounts of secondary BAs and indirectly altering BAs transporter activities. Female-predominant gut microbiome may have an increased capacity to generate secondary BAs (Figure 6). As BAs function as signaling molecules, variations in BA profiles resulting from sex-specific gut microbiomes could impact the activation of receptors and subsequently influence host health. It is worth noting that the gut microbiome composition in transplanted mice may differ from that in donor mice. To further explore this, additional experiments are needed to accurately identify bacterial compositions.

**Figure 6.**
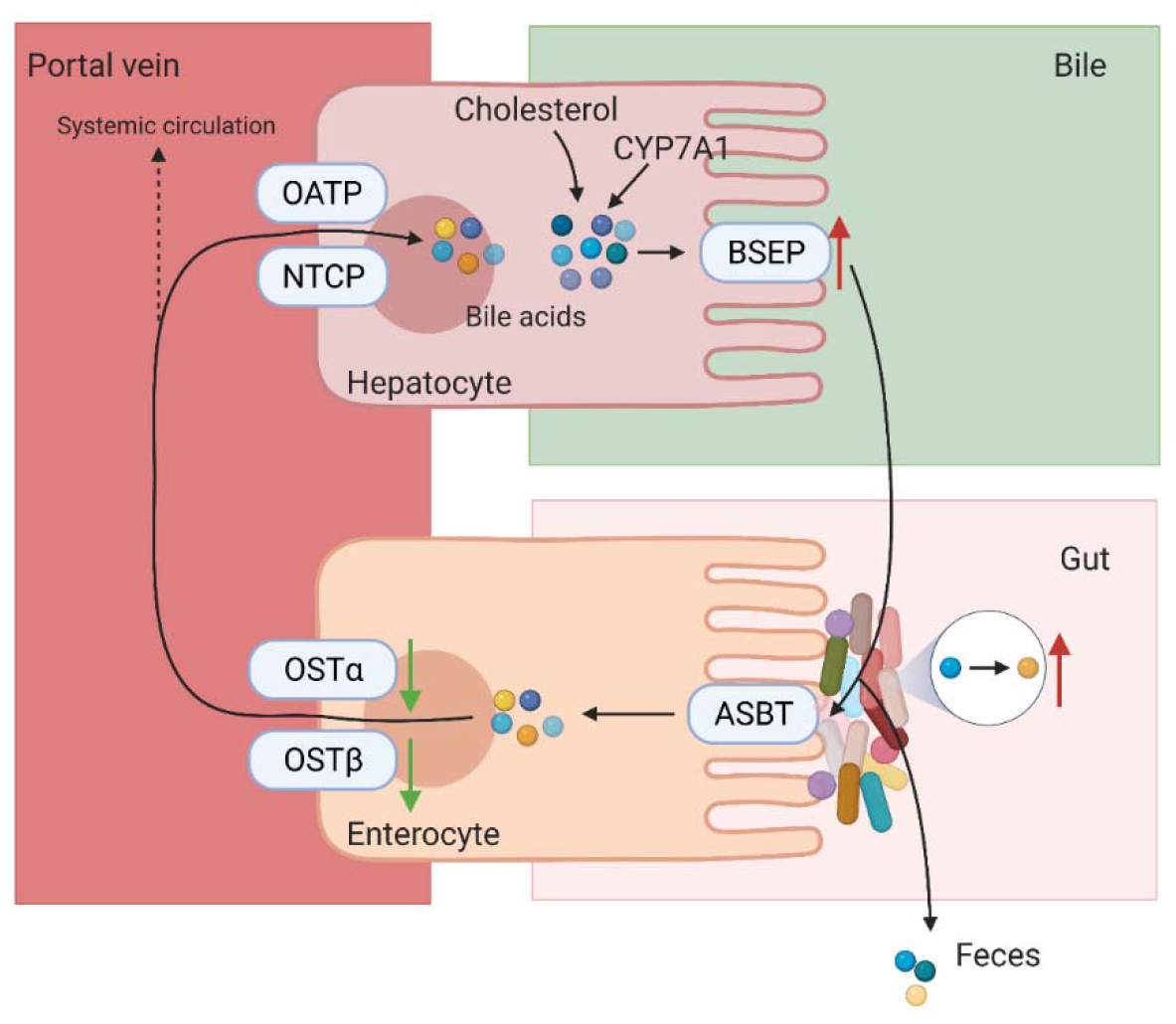
Schematic overview of BAs metabolism by sex-specific gut microbiome and host. BAs are synthesized in liver and undergo enterohepatic circulation with the help of transporters in liver and distal ileum. Sex-specific gut microbiome may influence BAs metabolism by directly producing more secondary BAs and indirectly changing BAs transporter activities, with female-predominant gut microbiome have the ability to produce more secondary BAs.

## Author Contributions

Supervision, funding acquisition and project administration, K.L.; conceptualization, Y.Y. and K.L.; methodology, Y.Y., C.L. and K.L.; writing—original draft preparation, Y.Y.; writing— review and editing, Y.Y., C.L., Y.H., H.R. and K.L. All authors have read and agreed to the published version of the manuscript.

## Supplementary Material

Supplementary Figure 1. Unconjugated/conjugated ratio were constant in two groups in feces, serum and liver. Expression of bile acids conjugation related gene in liver were observed no difference in two groups; Supplementary Figure 2. Initial and final body weight of recipient mice; Supplementary Table 1. Initial and final body weight of recipient mice; Supplementary Table 2. Fecal DNA concentration and purity after antibiotics treatment; Supplementary Table 3. Fecal DNA concentration and purity after fecal transplantation; Supplementary Table 4. MRM transi-tions of bile acids measured in the present study; Supplementary Table 5. Primers used in this study.

## Data Availability Statement

The datasets [GENERATED/ANALYZED] for this study can be found in the [NAME OF REPOSITORY] [LINK]. Please see the “Availability of data “ section of <u>Materials and data policies in the Author guidelines</u> for more detail

