## Supplemental material for "Sex-specific effects of gut microbiome on shaping bile acid metabolism"

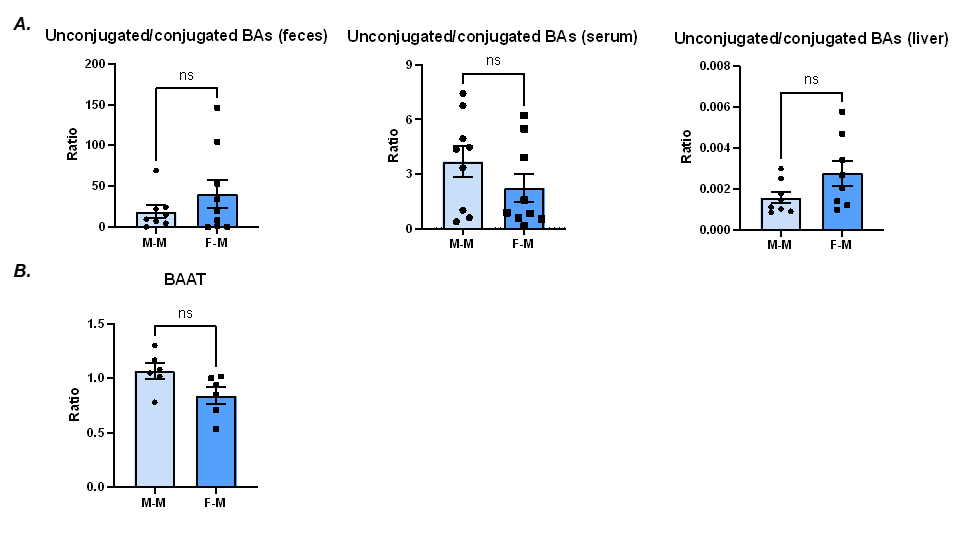

**Supplementary Figure 1.** Unconjugated/conjugated ratio were constant in two groups in feces, serum and liver. Expression of bile acids conjugation related gene in liver were observed no difference in two groups. (M-M: mice transplanted with male gut microbiome, F-M: mice transplanted with female gut microbiome; Data are expressed as mean ± SEM, unpaired t test, ns: no significant difference; *p<0.05, **p<0.01, ***p<0.001; n = 6 /group).

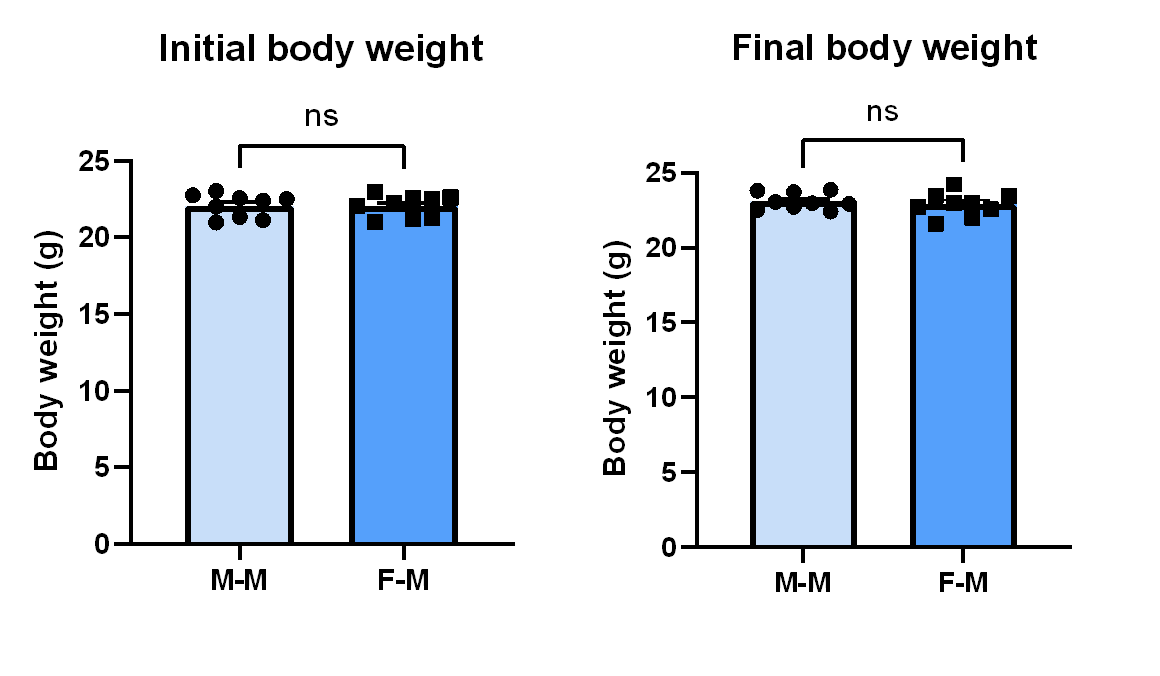

**Supplementary Figure 2.** Initial and final body weight of recipient mice. (M-M: mice transplanted with male gut microbiome, F-M: mice transplanted with female gut microbiome; Data are expressed as mean ± SEM, unpaired t test, ns: no significant difference; *p<0.05, **p<0.01, ***p<0.001; n = 9 /group).

**Supplementary Table 1.** Initial and final body weight of recipient mice

| Group | | Initial Body Weight (g) | Final Body Weight(g) |
| --- | --- | --- | --- |
| M-M(n=9) | 20.98 | | 22.72 |
|  | 21.13 | | 22.40 |
|  | 22.01 | | 23.71 |
|  | 23.03 | | 22.96 |
|  | 22.75 | | 22.93 |
|  | 22.40 | | 23.87 |
|  | 21.34 | | 22.50 |
|  | 22.58 | | 23.80 |
|  | 22.50 | | 23.05 |
| F-M(n=9) | 21.00 | | 23.02 |
|  | 22.97 | | 23.45 |
|  | 22.06 | | 24.22 |
|  | 21.20 | | 22.56 |
|  | 22.62 | | 23.00 |
|  | 22.26 | | 21.60 |
|  | 21.30 | | 22.01 |
|  | 22.58 | | 22.72 |
|  | 22.48 | | 23.46 |

**Supplementary Table 2.** Fecal DNA concentration and purity after antibiotics treatment

| Group | DNA concentration (ng/μL) | DNA purity (A260/280) |
| --- | --- | --- |
| M-M(n=9) | 5.3 | 1.62 |
|  | 7.7 | 1.54 |
|  | 3.6 | 1.42 |
|  | 3.7 | 1.48 |
|  | 4.5 | 1.48 |
|  | 4.3 | 1.42 |
|  | 2.9 | 1.27 |
|  | 3.4 | 1.63 |
|  | 4.6 | 1.43 |
| F-M(n=9) | 6.8 | 1.52 |
|  | 3.4 | 1.52 |
|  | 3.3 | 1.55 |
|  | 2.9 | 1.35 |
|  | 8.6 | 1.68 |
|  | 4.0 | 1.49 |
|  | 2.7 | 1.55 |
|  | 2.0 | 1.18 |
|  | 3.8 | 1.7 |

**Supplementary Table 3.** Fecal DNA concentration and purity after fecal transplantation

| Group | DNA concentration (ng/μL) | DNA purity (A260/280) |
| --- | --- | --- |
| M-M(n=9) | 26.4 | 1.81 |
|  | 34.7 | 1.82 |
|  | 30 | 1.83 |
|  | 37.2 | 1.83 |
|  | 22.5 | 1.76 |
|  | 169.5 | 1.82 |
|  | 37.8 | 1.78 |
|  | 40.9 | 1.82 |
|  | 29.2 | 1.79 |
| F-M(n=9) | 88.6 | 1.84 |
|  | 47.2 | 1.87 |
|  | 37.4 | 1.83 |
|  | 39.4 | 1.81 |
|  | 68.9 | 1.84 |
|  | 76.2 | 1.85 |
|  | 37.7 | 1.84 |
|  | 29.7 | 1.85 |
|  | 30.3 | 1.81 |

**Supplementary Table 4.** MRM transitions of bile acids measured in the present study

| Bile acids | abbre. | IS | | | mass | Q1 (m/z) | | Q3(m/z) | RT |
| --- | --- | --- | --- | --- | --- | --- | --- | --- | --- |
| Glycocholic acid hydrate | GCA | | d4-GCA | 465.62 | | | 464.3 | 74 | 11.53 |
| Glycohyocholic acid, sodium salt | GHCA | | d4-GCA | 487.6 | | | 464.3 | 74 | 9.61 |
| Glycodeoxycholic acid, sodium salt | GDCA | | d6-GDCA | 471.61 | | | 448.3 | 74 | 16.82 |
| Glycodehydrocholic acid, sodium salt | GDHCA | | d6-GDCA | 481.56 | | | 458.3 | 74 | 5.44 |
| Glycolithocholic acid | GLCA | | d4-GLCA | 433.62 | | | 432.3 | 74 | 18.31 |
| Glycochenodeoxycholic Acid | GCDCA | | d4-GUDCA | 471.6 | | | 448.3 | 74 | 18.34 |
| Glycohyodeoxycholic Acid | GHDCA | | d4-GUDCA | 449.6 | | | 448.3 | 74 | 11.49 |
| Glycoursodeoxycholic acid | GUDCA | | d4-GUDCA | 449.62 | | | 448.3 | 74 | 11.14 |
| Tauro α-muricholic acid sodium salt | TαMCA | | d4-TβMCA | 537.68 | | | 514.3 | 80/107 | 5.82 |
| Tauro β-muricholic acid sodium salt | TβMCA | | d4-TβMCA | 537.68 | | | 514.3 | 80/124 | 6.02 |
| Tauro ω-muricholic acid sodium salt | TωMCA | | d4-TβMCA | 537.68 | | | 514.3 | 80 | 5.44 |
| Taurohyocholic acid, sodium salt | THCA | | d4-CA | 537.69 | | | 514.3 | 80/107 | 8.02 |
| Taurocholic acid sodium salt hydrate | TCA | | d4-CA | 537.68 | | | 514.3 | 80/107 | 10.30 |
| Tauroursodeoxycholic Acid, Sodium Salt | TUDCA | | d4-TUDCA | 521.69 | | | 498.3 | 80 | 9.58 |
| Sodium taurochenodeoxycholate | TCDCA | | d9-TCDCA | 521.69 | | | 498.3 | 80/107 | 14.79 |
| Taurodehydrocholic acid | TDHCA | | d6-TDCA | 509.66 | | | 508.2 | 80 | 4.11 |
| Sodium taurodeoxycholate hydrate(xH2O) | TDCA | | d6-TDCA | 521.69 | | | 498.3 | 80 | 15.93 |
| Cholic acid | CA | | d4-CA | ‎408.579 | | | 407.3 | 407.3 | 15.73 |
| Hyocholic Acid | HCA | | d4-CA | 408.6 | | | 407.3 | 407.3 | 14.15 |
| α-muricholic acid | αMCA | | d4-CA | ‎408.579 | | | 407.3 | 407.3 | 11.71 |
| β-muricholic acid | βMCA | | d4-CA | ‎408.579 | | | 407.3 | 407.3 | 12.34 |
| ω-muricholic acid | ωMCA | | d4-CA | ‎408.579 | | | 407.3 | 407.3 | 11.12 |
| Ursodeoxycholic acid | UDCA | | d4-UDCA | 392.56 | | | 391.3 | 391.3 | 16.10 |
| Murodeoxycholic acid | MDCA | | d9-CDCA | 392.6 | | | 391.3 | 391.3 | 14.47 |
| Chenodeoxycholic acid | CDCA | | d9-CDCA | 392.56 | | | 391.3 | 391.3 | 18.04 |
| Dehydrocholic acid | DHCA | | d6-DCA | 402.52 | | | 401.2 | 401.2 | 9.60 |
| Deoxycholic acid | DCA | | d6-DCA | 392.56 | | | 391.3 | 391.3 | 18.21 |
| Lithocholic acid | LCA | | d4-LCA | 376.5726 | | | 375.3 | 375.3 | 19.42 |

**Supplementary Table 5.** Primers used in this study

| Gene name | Forward Primer Sequence (5’ to 3’) | Reverse Primer Sequence (5’ to 3’) | Tm (℃) |
| --- | --- | --- | --- |
| β-Actin | CGTGCGTGACATCAAAGAGAA | TGGATGCCACAGGATTCCAT | 55 |
| Cyp7a1 | AGCAACTAAACAACCTGCCAGTACT | GTCCGGATATTCAAGGATGCA | 54 |
| Bsep | CTGCCAAGGATGCTAATGCA | CGATGGCTACCCTTTGCTTCT | 55 |
| Oatp | CAGTCTTACGAGTGTGCTCCAGAT | ATGAGGAATACTGCCTCTGAAGTG | 55 |
| Ntcp | ATGACCACCTGCTCCAGCTT | GCCTTTGTAGGGCACCTTGT | 55 |
| Asbt | TGGGTTTCTTCCTGGCTAGACT | TGTTCTGCATTCCAGTTTCCAA | 55 |
| Ostα | TGTTCCAGGTGCTTGTCATCC | CCACTGTTAGCCAAGATGGAGAA | 55 |
| Ostβ | GATGCGGCTCCTTGGAATTA | GGAGGAACATGCTTGTCATGAC | 55 |
